## Supplementary Information for "AMPK deficiency in smooth muscles causes persistent pulmonary hypertension after birth and premature death"

### Supplemental Methods

**Western Blot** Liver, kidney and ventricle were lysed by sonication. 33µg of each lysate was run on gels. AMPK subunit protein expression was analyzed using precast 4-12% BisTris gels in MOPS buffer. Proteins were transferred to nitrocellulose membranes using an Xcell II blot module and probed with antibodies against AMPK subunits. AMPK-α1 (ab3759, cell signalling) and AMPK-α1 (ab105028, cell signalling) antibodies were Abcam. GAPDH antibody (2118S) was from cell signalling.

**TASK-1 expressing HEK293 cells** were generated as follows. Full length cDNAs encoding human TASK-1 (hTASK-1) channels, a kind gift from Dr. S. A. N. Goldstein (Department of Pediatrics and Institute for Molecular Pediatric Sciences, Pritzker School of Medicine, University of Chicago), were originally subcloned into the mammalian expression vectors pRAT and pMAX(+) via Bgl II/Sal I and Xba I/Xma I restriction site combinations respectively. To generate HEK293 cell lines stably expressing either wild-type hTASK-1, wild-type hTASK-3 or the hTASK-3(S55A) mutant, cells were transfected with either pRAT/hTASK-1, pMAX/hTASK-3 or pXX-hTASK-3(S55A) constructs respectively, using the PolyFect transfection reagent (Qiagen, Hybaid Ltd, Teddington, UK) according to manufacturer's instructions. Stable HEK293 cell lines were achieved by antibiotic selection with G-418 (1mg/ml, Gibco-BRL, Paisley, UK) added to the medium 3 days after transfection. Selection was applied for 4 weeks (media changed every 4-5 days), after which time individual colonies were picked and seeded in T25 flasks and allowed to reach confluence. They were then transferred to T75 flasks for further culture and electrophysiological screening. Cells were harvested from culture flasks by trypsinization and plated onto coverslips 24-48h before use in electrophysiological studies. Transfection of hTASK-1 and hTASK-3 channels was considered successful if the currents elicited by the whole-cell voltage ramp protocol (see below) were: a) significantly (> 4-fold) larger than untransfected HEK293 cell $K^+$ currents; b) reliably described by the Goldman Hodgkin and Katz (GHK) equation and; c) activated by pH 8.4 and inhibited by pH 6.4 (and the pH-sensitive currents demonstrated GHK rectification). In addition, hTASK1 was insensitive to ruthenium red, while hTASK3 was sensitive as previously described (Czirjak and Enydei Mol Pharm. 2003; 63: 646-652; DOI: https://doi.org/10.1124/mol.63.3.646). Whole-cell patch-clamp recordings were recorded from HEK293 cells stably expressing wild-type hTASK-1. Coverslip fragments with attached cells were transferred to a continuously perfused recording chamber (perfusion rate 3-5 ml/min, volume ca 200µl) mounted on the stage of an inverted microscope. Cells were perfused with a solution containing (in mM): 135 NaCl, 5 KCl, 1.2 $MgCl_2$, 5 HEPES, 2.5 $CaCl_2$, 10 D-glucose (pH 7.4 with KOH). Patch electrodes (resistance 4-7MΩ) were filled with intracellular solution consisting of (in mM): 10 NaCl, 117 KCl, 2 $MgCl_2$, 11 HEPES, 11 EGTA, 1 $CaCl_2$, 2 $Na_2ATP$ (pH 7.2 with KOH). All chemicals were obtained from Sigma-Aldrich (Poole, UK). Outward $K^+$ currents were recorded at 37°C (unless otherwise stated) using standard ramp protocols; cells were voltage-clamped at a holding potential of –70mV, then stepped to -100mV and a voltage ramp (500ms duration) immediately applied from -100 mV to +60mV. Data were acquired and digitized via a Digidata 1322A in combination with an Axopatch 200B amplifier and Clampex 9 software (Molecular Devices, Foster City, CA). Currents were sampled at 2kHz and low-pass filtered at 1kHz. Offline data analysis was conducted using Clampfit 9 software (Molecular Devices, Foster City, CA.) The effects of drugs were determined using the paired Student's *t-test* on the pre and post-application of drugs at the indicated concentrations. Differences were considered significant when $P<0.05$. All values stated are as mean ± SEM.

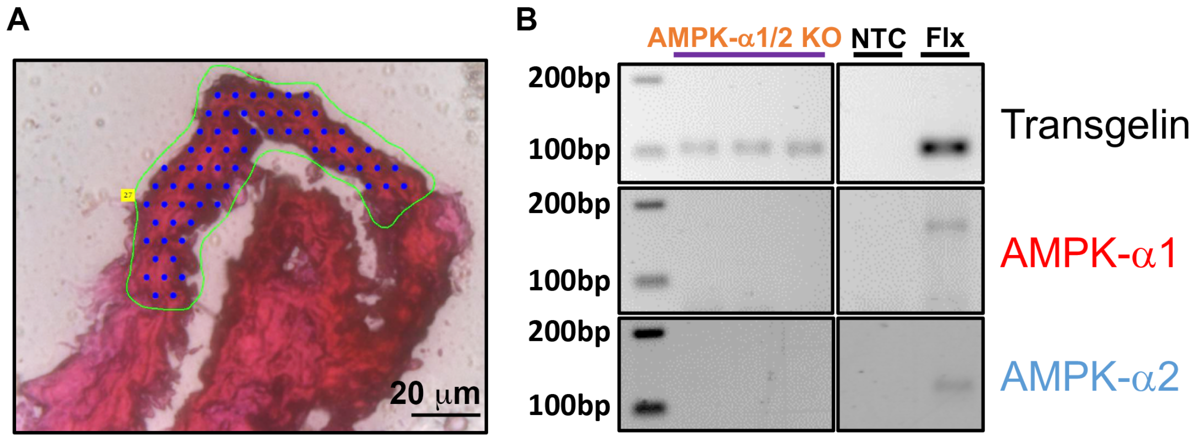


**Supplementary Figure 1. Laser microdissection and end-point PCR confirms AMPK-α1/α2 deletion in pulmonary arterial smooth muscles.** A, Exemplar image shows region of interest (green line) at x40 magnification that were used to direct laser microdissection of the medial layer of pulmonary arterial section. The blue dots show where the energy pulses were targeted that lift the sample off the slide and transfer it into an adhesive cap placed directly above the sample; Scale bar 20μm. B, Gel shows single cell end-point RT-PCR amplicons for transgelin, AMPK-α1 and AMPK-α2 from AMPK-α1/α2 knockout and AMPK-α1/α2 floxed (Flx) mice. NTC = negative control (eluate from the RNA extraction for which no sample material was added).

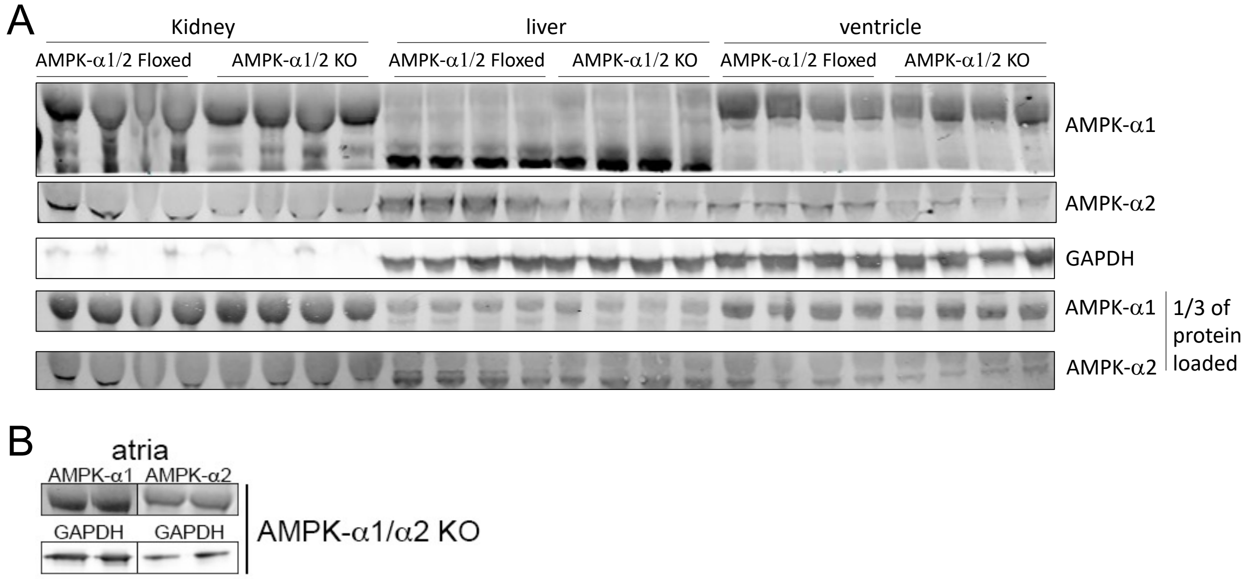


**Supplementary Figure 2. Western blot reveals no loss of AMPK-α1 or AMPK-α2 expression in cardiac ventricles following AMPK-α1/α2 deletion by transgelin-Cre.** A, Exemplar western blots for AMPK-α1 and AMPK-α2 protein levels in kidney, liver and left ventricle of the heart for AMPK-α1/α2 Floxed and AMPK-α1/α2 knockout (KO) mice, relative to loading control (GAPDH). B, Exemplar western blots for AMPK-α1 and AMPK-α2 protein levels in the atria of AMPK-α1/α2 knockout (KO) mice, relative to loading control (GAPDH).

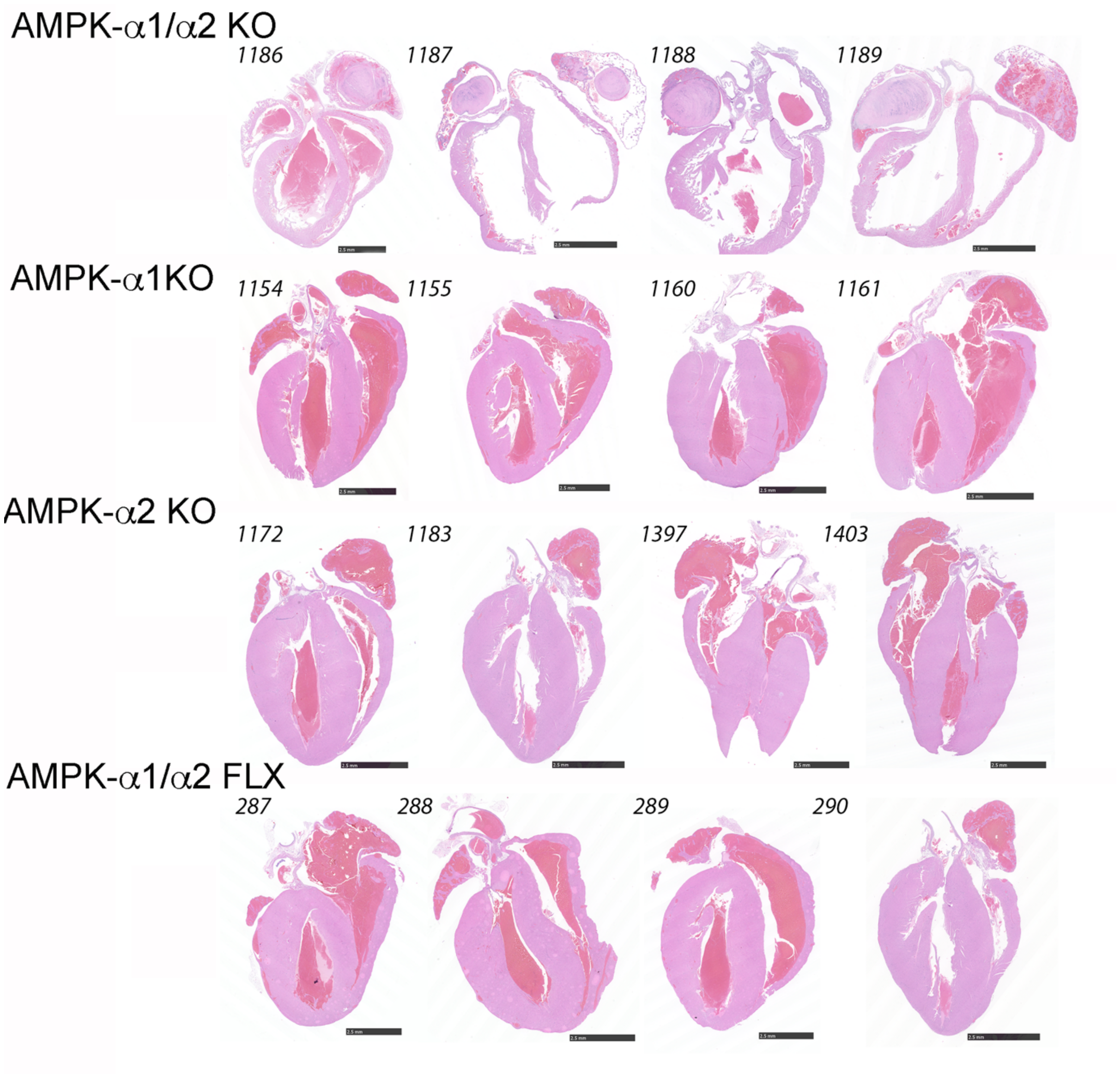


**Supplementary Figure 3. Remodelling of the heart in terminal samples from AMPK-α1/α2 knockouts.** Sub-gross images of heart slices from terminal samples stained with Hematoxylin-Eosin for: AMPK-α1/α2 knockouts (AMPK-α1/α2 KO); AMPK-α1 knockout (AMPK-α1 KO); AMPK-α2 knockout (AMPK-α2 KO); AMPK-α1/α2 floxed (AMPK-α1/α2 FLX). Scale bars 2.5mm.

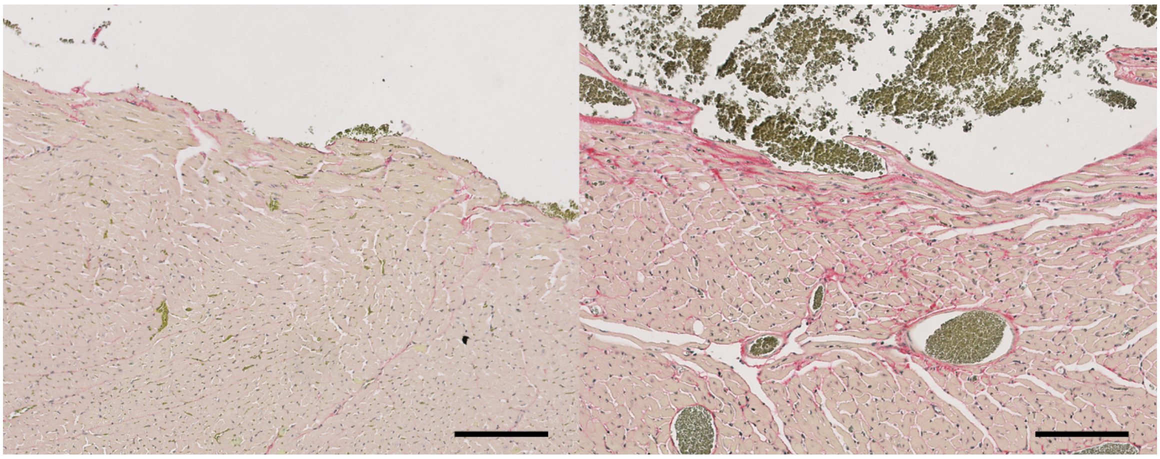

**Supplementary Figure 4. AMPK-α1/α2 deletion precipitates fibrosis of the left ventricle.** Exemplar images of the left ventricle from age-matched AMPK-α1/α2 FLX (left) and AMPK-α1/α2 knockout (right) mice shows relative Picro-Sirius Red staining (red; erythrocytes are stained green (not analysed), counterstain is pale yellow. Scale bars 100 μm.

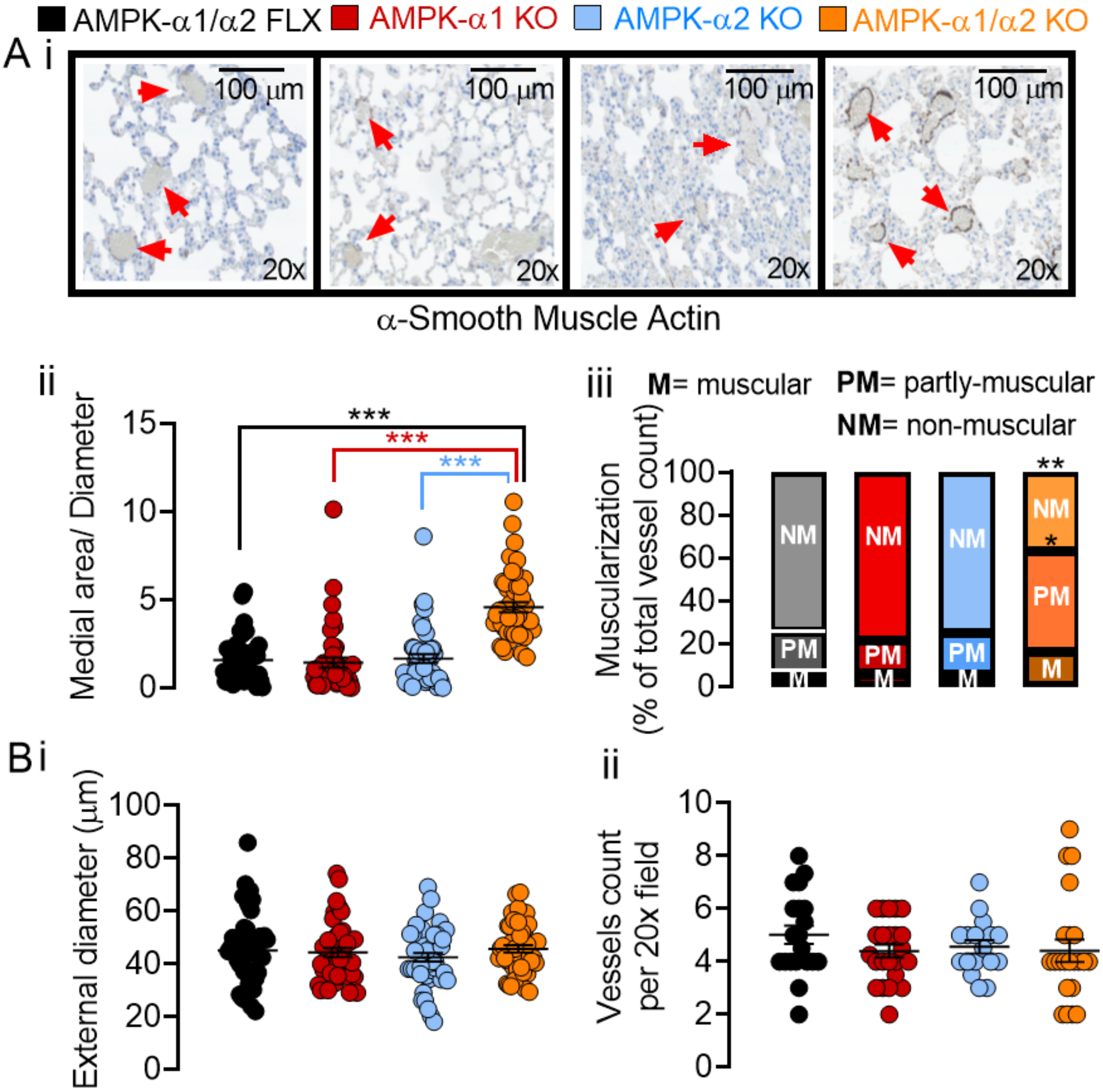


**Supplementary Figure 5. Analysis of pulmonary arterial remodelling triggered by AMPK-α1/α2 deletion by total vessel counts rather than by mouse.** Ai, Representative images of lung slices from terminal samples stained for α-smooth muscle actin, together with scatter plots of (ii) medial area corrected by diameter and (iii) degree of muscularization for all pulmonary arteries analyzed for AMPK-α1/α2 KOs after death at 7-10 weeks, and age-matched AMPK- α1/α2 floxed, AMPK-α1 KOs and AMPK-α2 KOs. B, Scatter plots show the mean ± SEM for the (i) external diameter and (ii) number of vessels found per 20x field for the analysis shown in B (n = 122-140 arteries, 7 fields, 4 mice). *P<0.05, **P<0.01, ***P<0.001.

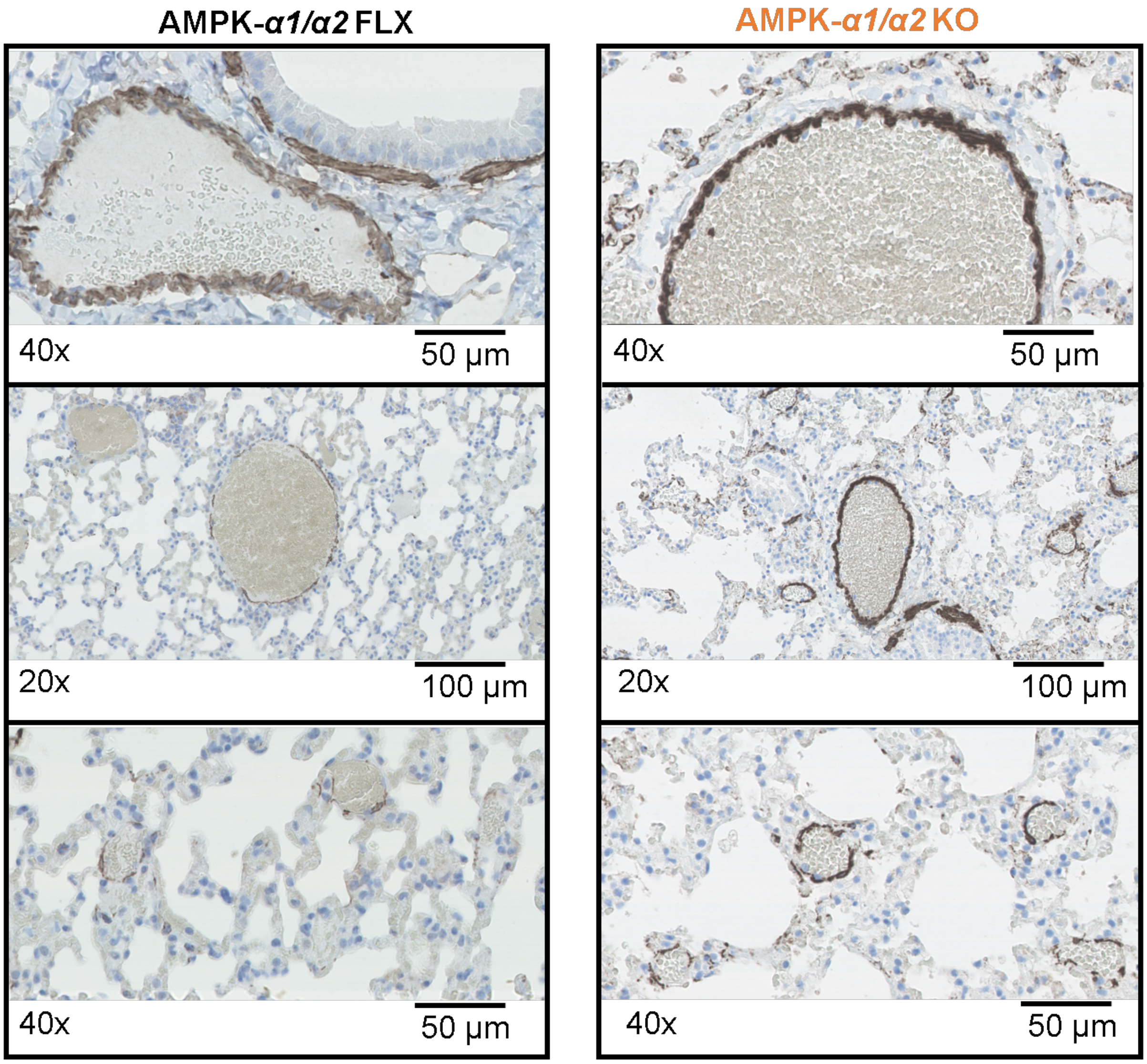


**Supplementary Figure 6. Remodelling of pulmonary arterial media in AMPK-α1/α2 knockouts.** Representative high-resolution images of pulmonary arteries in terminal lung slices of AMPK-α1/α2 FLX (left) and AMPK-α1/α2 knockout (right) mice stained for α-smooth muscle actin.

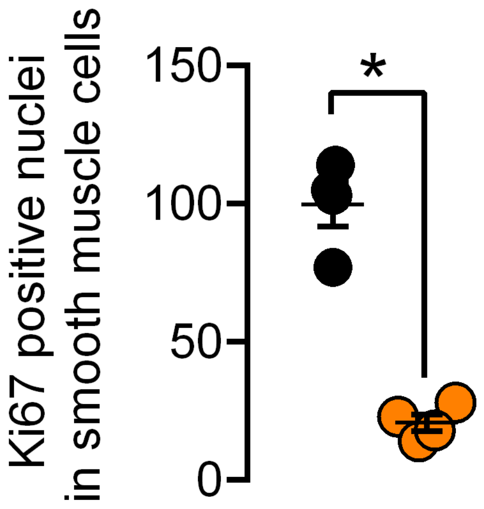


**Supplementary Figure 7. Reduced Ki67 positive nuclei in pulmonary arterial myocytes of AMPK-α1/α2 knockouts after birth.** Scatter plot shows the mean ± SEM for Ki67 labelled nuclei in smooth muscle cells of pulmonary arteries in-situ within lung slices from AMPK-α1/α2 FLX (black, n = 4 mice) and AMPK-α1/α2 KO (orange, n=4 mice). * $P<0.05$ vs AMPK-α1/α2 FLX.

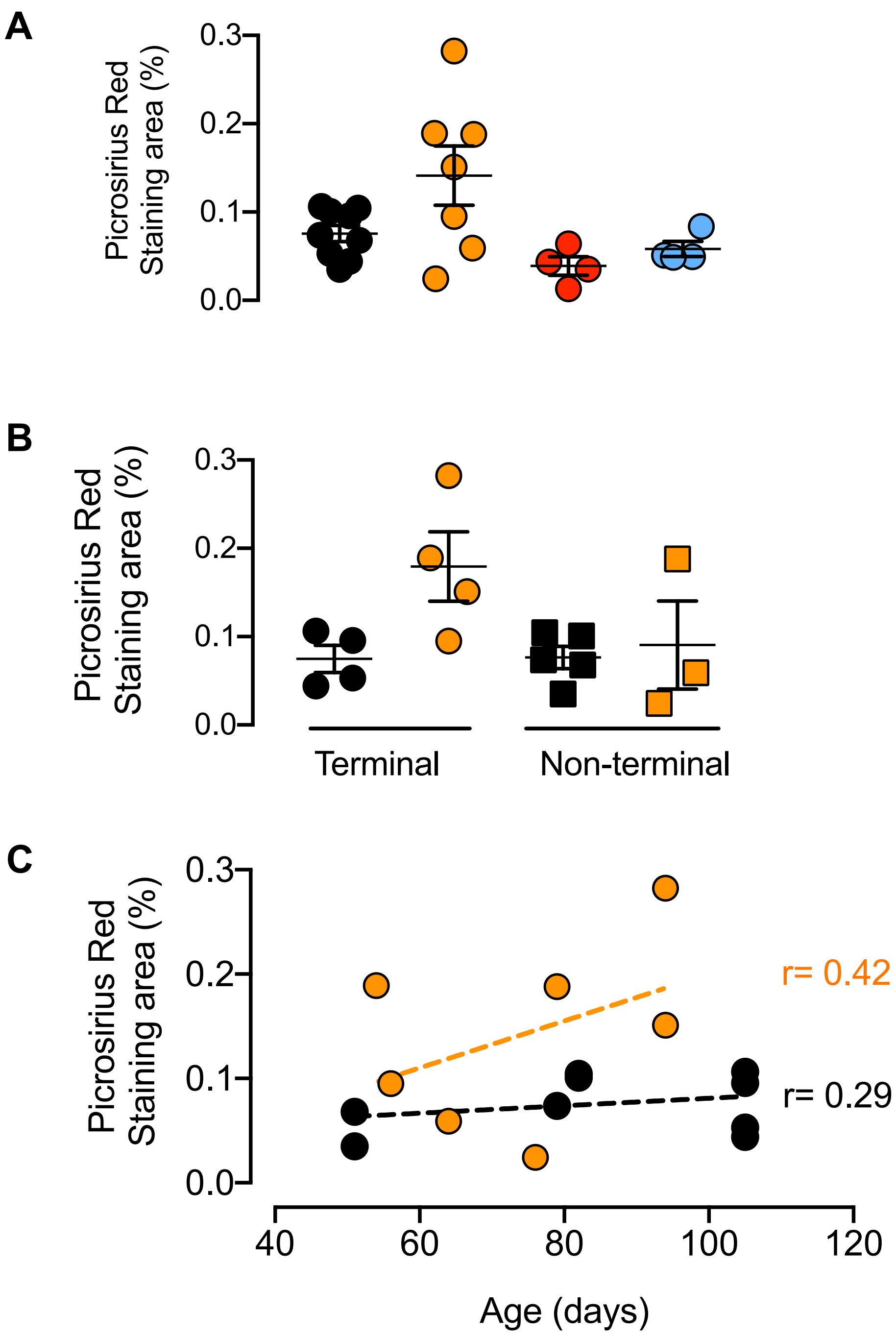


**Supplementary Figure 8. Parenchymal fibrosis is not triggered by AMPK-α1/α2 deletion.** A, Scatter plots show the mean ± SEM for analysis of Picro-Sirius Red staining by Kruskall-Wallis test. AMPK-α1/α2 FLX (black, n = 8 mice) and AMPK-α1/α2 KO (orange, n = 7 mice), AMPK-α1 KO (red, n = 4 mice), AMPK-α2 KO (blue, n = 4 mice). B, as in A but for comparison of terminal and non-terminal samples from AMPK-α1/α2 KO vs AMPK-α1/α2 FLX. C, linear regression analysis of Picro-Sirius Red staining versus age.

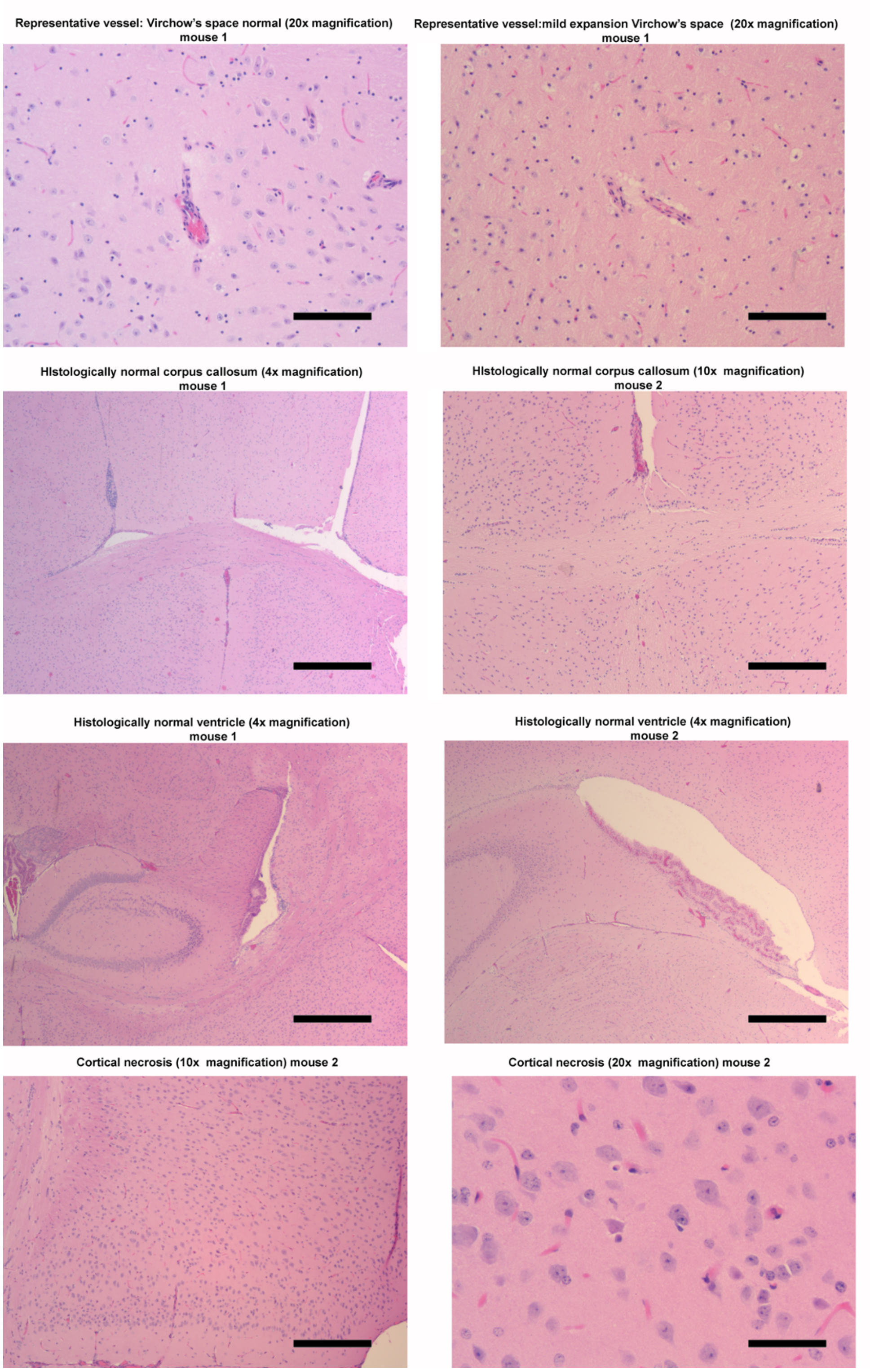


**Supplementary Figure 9. Histology on terminal AMPK-α1/α2 knockouts indicates no lesion of the brain.** Panels, from top to bottom, show representative images of Virchow's space, corpus callosum, IVth ventricle, and cortical necrosis observed in one mouse of three mice studied (likely caused by cortical ischaemia at time of death). In two mice mild expansion of the Virchow's space was observed, that was likely caused by a fixation/processing artefact rather than oedema because proteinaceous fluid is absent. The scale bars are: 4x magnification, 50μm; 10x magnification, 120μm; 20x magnification, 300μm.

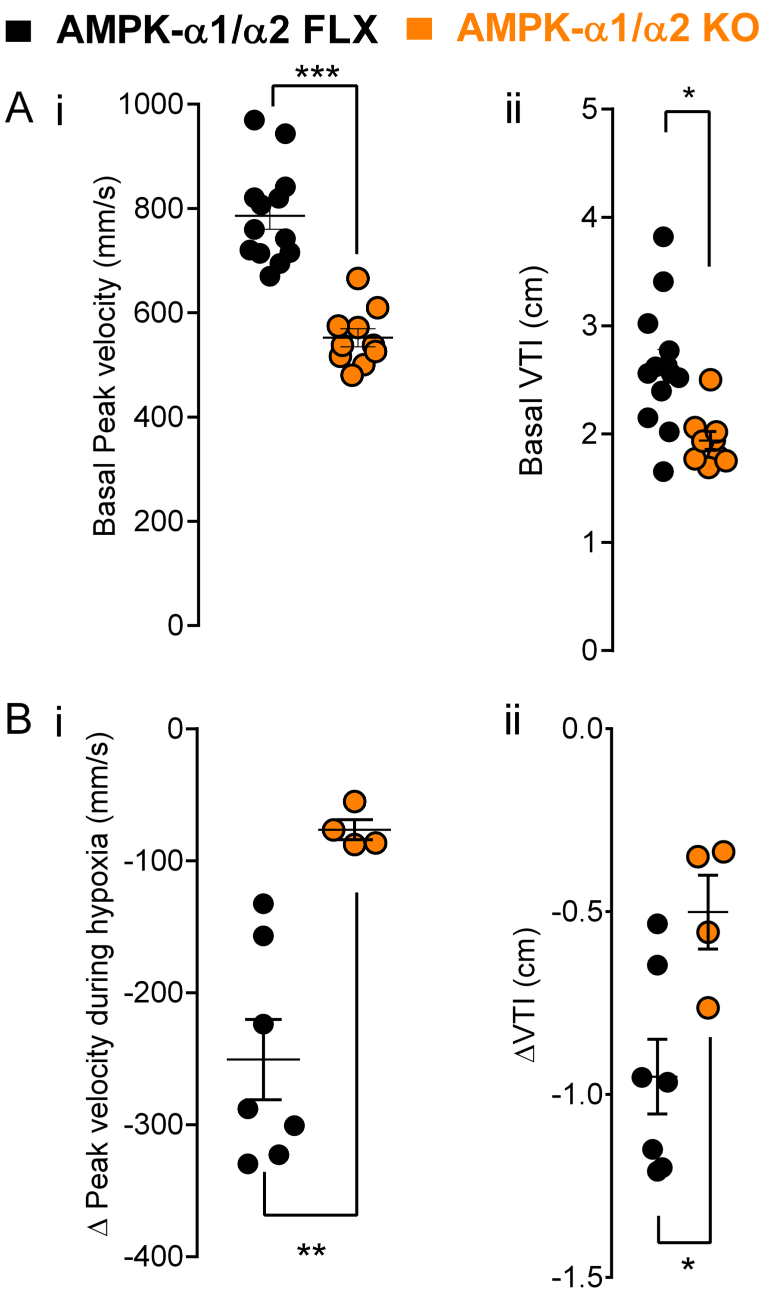


**Supplementary Figure 10. Reduced normoxic pulmonary flow and impaired hypoxic pulmonary vasoconstriction in AMPK-α1/α2 identified by parallel changes in peak velocity and velocity time integral.** A, Scatter plots show the mean ± SEM for normoxic peak velocity (i) and normoxic velocity time integral (VTI) (ii). (B) Scatter plots for maximum change in peak velocity (i) and VTI (ii) observed during 8% $O_2$. Data from AMPK-α1/α2 floxed (AMPK-α1/α2 FLX) and AMPK-α1/α2 knockouts (AMPK-α1/α2 KO, n=4-9). * P<0.05, ** P<0.01, *** P<0.001 vs AMPK-α1/α2 FLX.

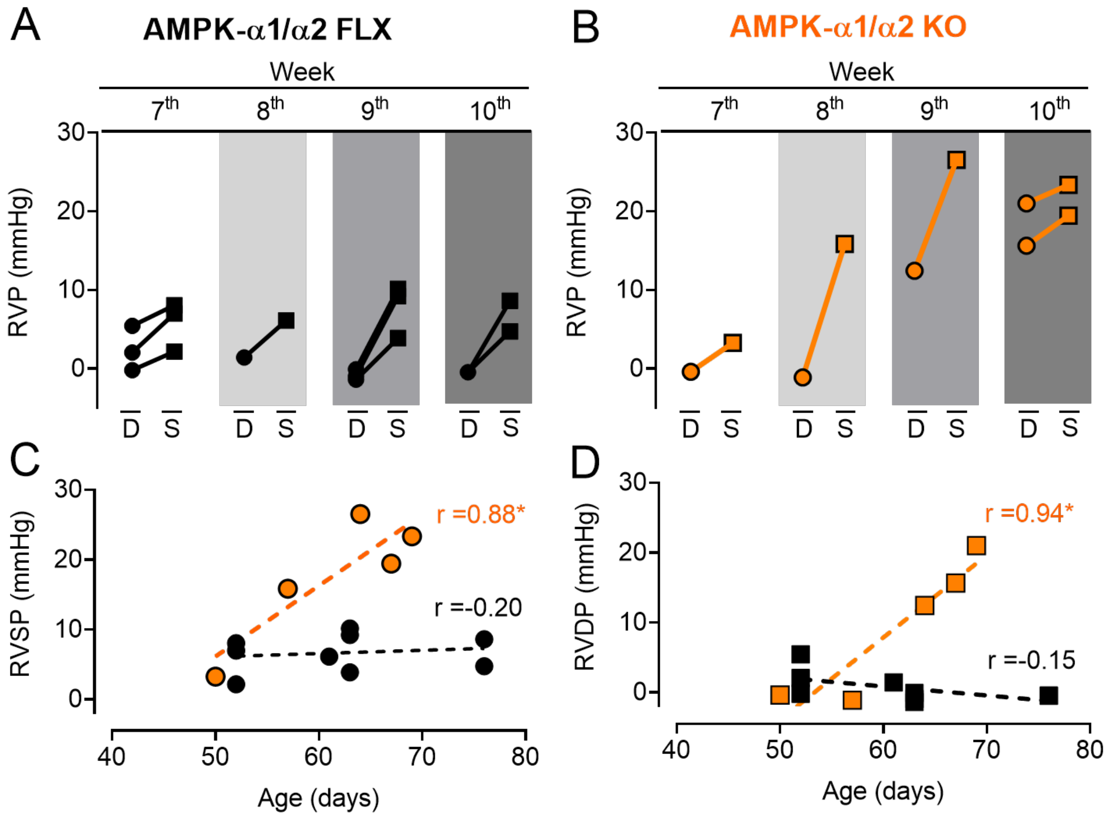


**Supplementary Figure 11. Right ventricular systolic pressure of AMPK-α1/α2 knockouts increases rapidly with age.** Graphs illustrate age-dependent changes in right ventricular systolic (S) and diastolic (D) pressures for (A) AMPK-α1/α2 floxed (AMPK-α1/α2 FLX) and (B) AMPK-α1/α2 knockout (AMPK-α1/α2 KO) mice. Scatter plots show values of (C) right ventricular systolic and (D) diastolic pressures against age for AMPK-α1/α2 FLX and AMPK-α1/α2 KO (n=5-9). Linear regression analysis indicates (coefficient of determination $r^2$ = 0.77) that right ventricular systolic pressure of AMPK-α1/α2 knockouts (n = 5) increases rapidly with age between 50 and 70 days (orange), while for age-matched controls (AMPK-α1/α2 floxed, n = 9) there was no correlation between right ventricular systolic pressure and age (black). * P<0.05.

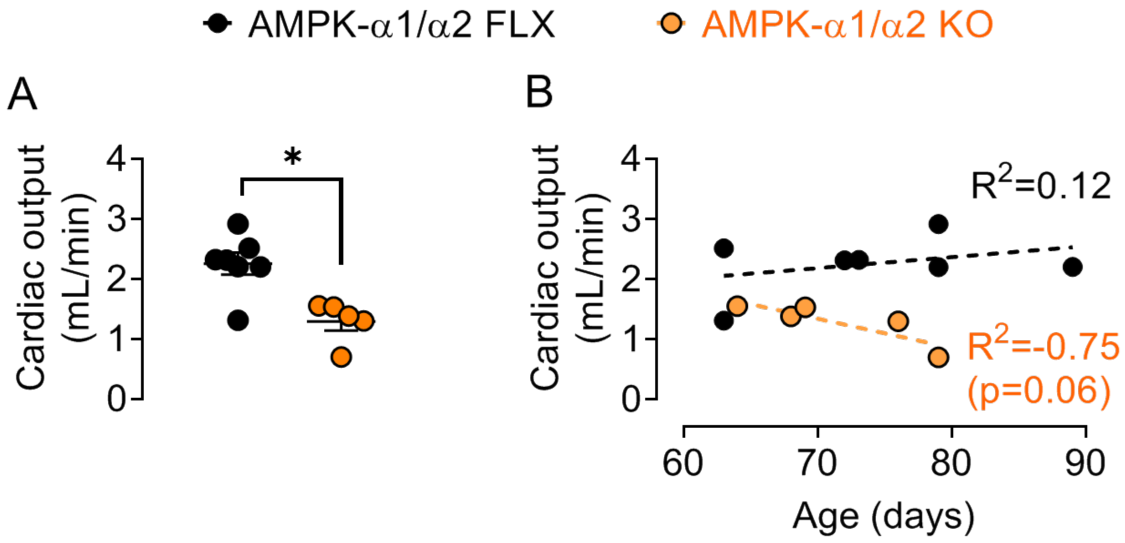


**Supplementary Figure 12. Right ventricular cardiac output of AMPK-α1/α2 knockouts decreases with age.** Scatter plots show (A) mean ± SEM for right ventricular cardiac output and (B) age-dependent changes in right ventricular cardiac output for AMPK-α1/α2 floxed (AMPK-α1/α2 FLX) and AMPK-α1/α2 knockout (AMPK-α1/α2 KO) mice (n=5-9; * P<0.05). Linear regression analysis indicates (coefficient of determination $R^2$ = -0.75) that right ventricular cardiac output of AMPK-α1/α2 knockouts (n = 5) decreases with age between 50 and 70 days (orange), while for age-matched controls (AMPK-α1/α2 floxed, n = 9) there was no correlation between cardiac output and age between 50 and 70 days (black). Cardiac output was derived from the heart rate and flow volume calculated by the velocity-time integral (VTI) and cross-sectional area (CSA) of the valve: CO [L/min] = HR x VTI x CSA (cross sectional area of murine pulmonary artery; diameter = 0.44 ± 0.06[1]; $A=\pi r^2$).

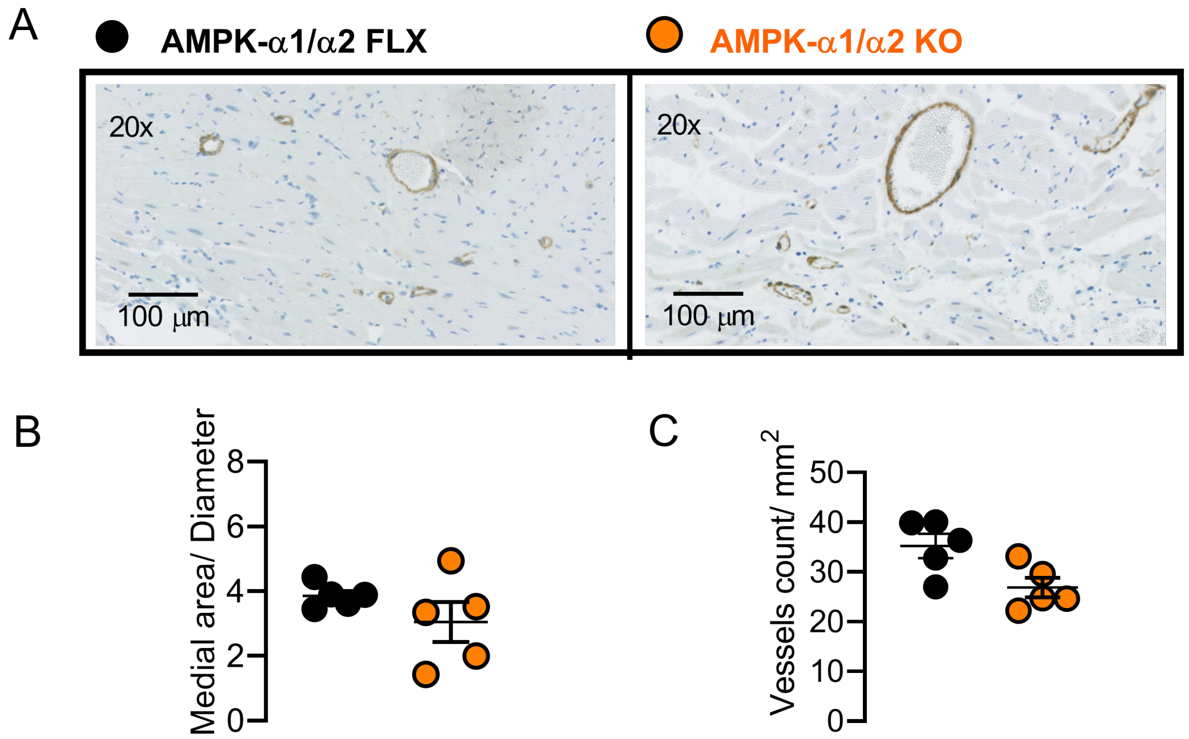


**Supplementary Figure 13. AMPK-α1/α2 deletion driven by Cre expression via the transgelin promoter does not affect right ventricular blood vessel number or medial thickness**. A, Representative images of right ventricles from samples stained for α-smooth muscle actin. Scatter plot shows measurements of (B) medial area/diameter (n = 5) and (C) vessel count/mm$^2$ from AMPK-α1/α2 KOs vs age-matched AMPK-α1/α2 floxed controls (n=5; Note. Non-parametric Mann-Whithney for C is p=0.056).

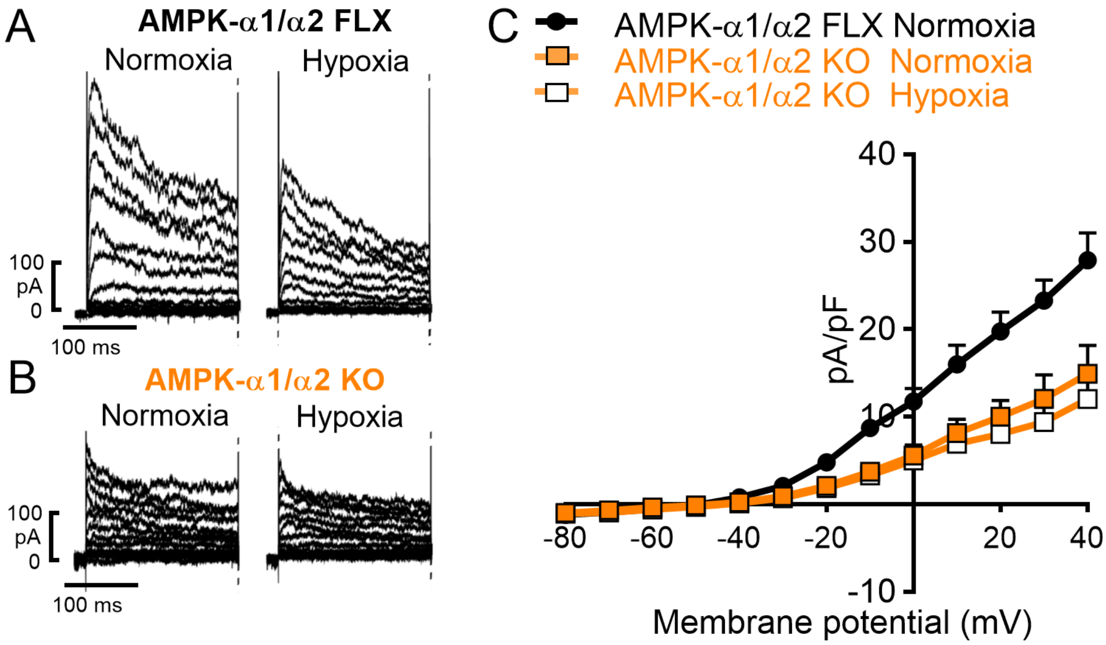


**Supplementary Figure 14. AMPK-α1/α2 deletion reduces $K_V$1.5 current density and blocks $K_V$ inhibition by hypoxia**. A-B, Panels show example records for $K_V$ currents recorded during control conditions and following extracellular during hypoxia (~6% $O_2$) in acutely isolated pulmonary arterial myocytes from either (A) AMPK-α1/α2 floxed (AMPK-α1/α2 FLX) or (B) AMPK-α1/α2 knockouts (AMPK-α1/α2 KO). C, Comparison of current-voltage relationship for $K_V$ currents recorded in pulmonary arterial myocytes from AMPK-α1/α2 floxed and AMPK-α1/α2 KO under control conditions, and the effect of hypoxia the current-voltage relationship of KV currents in pulmonary arterial myocytes from AMPK-α1/α2 KOs (n = 5 cells from at least 3 mice).

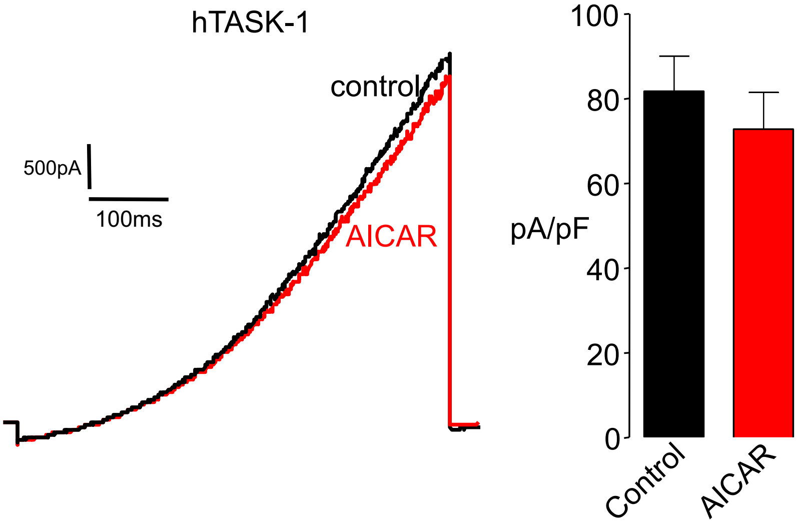


**Supplementary Figure 15**. **AMPK activation does not regulate TASK-1**. Lefthand Panel shows example records for $K^+$ currents in HEK 293 cells transfected with human KCNK3 (hTASK1) under control conditions and after pre-incubation (1hr) with AICAR (1mM). Right hand panel shows bar chart comparing mean ± SEM for the peak TASK1 current recorded under control conditions (n = 4) with that recorded in the presence of AICAR (1mM; n = 4).

Right ventricle

| | AMPK-α1/α2 FLX | | | | AMPK-α1 KO | | | | AMPK-α2 KO | | | | AMPK-α1/α2 KO | | | |
|---|---|---|---|---|---|---|---|---|---|---|---|---|---|---|---|---|
| | Mean | | Sem | n | Mean | | Sem | n | Mean | | Sem | n | Mean | | Sem | n |
| Systolic Volume (μL) | 2.2 | ± | 0.9 | (4) | 1.3 | ± | 0 | (4) | 2.6 | ± | 0.9 | (4) | 7.4 | ± | 1.5 | (5)* |
| Diastolic Volume (μL) | 9 | ± | 2.6 | (4) | 6.5 | ± | 0.7 | (4) | 7.1 | ± | 1.1 | (4) | 12.4 | ± | 2.4 | (5) |
| Stroke Volume (μL) | 6.8 | ± | 1.8 | (4) | 5.2 | ± | 0.6 | (4) | 4.5 | ± | 0.4 | (4) | 5.1 | ± | 1.9 | (5) |
| Ejection Fraction (%) | 78.4 | ± | 4.4 | (4) | 79.4 | ± | 1.6 | (4) | 66.4 | ± | 7.8 | (4) | 38.3 | ± | 10.6 | (5)** |
| Fractional Shortening (%) | 44.7 | ± | 4.3 | (4) | 44.8 | ± | 1.7 | (4) | 35 | ± | 5.8 | (4) | 17.9 | ± | 5.5 | (5) |

Left ventricle

| | AMPK-α1/α2 FLX | | | | AMPK-α1 KO | | | | AMPK-α2 KO | | | | AMPK-α1/α2 KO | | | |
|---|---|---|---|---|---|---|---|---|---|---|---|---|---|---|---|---|
| | Mean | | Sem | n | Mean | | Sem | n | Mean | | Sem | n | Mean | | Sem | n |
| Systolic Volume (μL) | 20.8 | ± | 3.1 | (4) | 16.2 | ± | 4 | (4) | 22.6 | ± | 4.3 | (4) | 105.7 | ± | 17.4 | (5)* |
| Diastolic Volume (μL) | 72.7 | ± | 3.7 | (4) | 61.2 | ± | 7.7 | (4) | 62.8 | ± | 5.1 | (4) | 129.8 | ± | 17.6 | (5)* |
| Stroke Volume (μL) | 51.9 | ± | 1.8 | (4) | 45.1 | ± | 6.4 | (4) | 40.2 | ± | 1.1 | (4)* | 24.2 | ± | 3.2 | (5)** |
| Ejection Fraction (%) | 71.8 | ± | 3.1 | (4) | 73.9 | ± | 5.2 | (4) | 65 | ± | 4.1 | (4) | 20 | ± | 3.7 | (5)** |
| Fractional Shortening (%) | 40.8 | ± | 2.6 | (4) | 42.6 | ± | 4.4 | (4) | 35.3 | ± | 3.1 | (4) | 9.2 | ± | 1.8 | (5)** |
| Cardiac Output (ml/min) | 28.5 | ± | 0.9 | (4) | 25.4 | ± | 2.9 | (4) | 21.1 | ± | 0.3 | (4)* | 11.8 | ± | 1.9 | (5)** |
| Heart rate (bpm) | 569.1 | ± | 9.1 | (4) | 576.7 | ± | 8.5 | (4) | 541.2 | ± | 13.4 | (4) | 508.4 | ± | 10.2 | (5)** |

Pulmonary flow

| | AMPK-α1/α2 FLX | | | | AMPK-α1 KO | | | | AMPK-α2 KO | | | | AMPK-α1/α2 KO | | | |
|---|---|---|---|---|---|---|---|---|---|---|---|---|---|---|---|---|
| | Mean | | Sem | n | Mean | | Sem | n | Mean | | Sem | n | Mean | | Sem | n |
| Peak velocity (mm/s) | 774.4 | ± | 38.5 | (6) | 737.5 | ± | 38.3 | (4) | 721.7 | ± | 20.2 | (7) | 531.6 | ± | 15.35 | (4)** |
| VTI (cm) | 2.7 | ± | 0.3 | (6) | 2.5 | ± | 0.1 | (4) | 2.7 | ± | 0.1 | (7) | 1.7 | ± | 0.1 | (4)** |

**Supplementary Table I. AMPK-α1/α2 deletion causes right and left ventricular myopathy and dysfunction.** Doppler ultrasound parameters for right and left ventricular function for AMPK-α1/α2 floxed (AMPK-α1/α2 FLX), AMPK-α1 knockouts (AMPK-α1 KO), AMPK-α2 knockouts (AMPK-α2 KO) and AMPK-α1/α2 knockouts (AMPK-α1/α2 KO). n=4-6, * P<0.05, ** P<0.01, *** P<0.001.

| AMPK-α1/α2 KO | Lateral ventricles | Corpus callosum/ Striatum/ Cortex area | Cortical neuronal necrosis | Enlargement of Virchow's space |
|---|---|---|---|---|
| 1 | Normal | Normal | No | Mild |
| 2 | Normal | Normal | Yes | No |
| 3 | Normal | Normal | No | Mild |

**Supplementary Table II. Qualitative histological assessment of brain sections of terminal AMPK-α1/α2 knockouts reveals no central pathology.** Qualitative histological assessment (exemplar images in **Figure E7**) of surface area identifies no evidence of ventricular expansion or reduction of striatal, cortical and corpus callosum volumes in section of brains from n =3 AMPK-α1/α2 knockouts. In two of the mice mild expansion of the Virchow's space was observed, that is likely caused by a fixation/processing artefact rather than edema because proteinaceous fluid is absent. One mouse exhibited signs of cortical neuronal necrosis, most likely caused by cortical ischaemia secondary to cardiac failure.
